# Computational dissociation of dopaminergic and cholinergic effects on action selection and inhibitory control

**DOI:** 10.1101/645093

**Authors:** Eduardo A. Aponte, Dario Schöbi, Klaas E. Stephan, Jakob Heinzle

**Affiliations:** Translational Neuromodeling Unit, Institute for Biomedical Engineering, University of Zurich & ETH Zurich, Wilfriedstrasse 6, 8002, Zurich, Switzerland; Wellcome Centre for Human Neuroimaging, University College London. 12 Queen Square, London, WC1N, 3AR, UK; Max Planck Institute for Metabolism Research, Gleueler Strasse 50, 50931 Cologne, Germany

## Abstract

**Background:** Patients with schizophrenia make more errors than healthy subjects on the antisaccade task. In this paradigm, participants are required to inhibit a reflexive saccade to a target and to select the correct action (a saccade in the opposite direction). While the precise origin of this deficit is not clear, it has been connected to aberrant dopaminergic and cholinergic neuromodulation.

**Methods:** To study the impact of dopamine and acetylcholine on inhibitory control and action selection, we administered two selective drugs (levodopa 200mg/galantamine 8mg) to healthy volunteers (N=100) performing the antisaccade task. A computational model (SERIA) was employed to separate the contribution of inhibitory control and action selection to empirical reaction times and error rates.

**Results:** Modeling suggested that levodopa improved action selection (at the cost of increased reaction times) but did not have a significant effect on inhibitory control. By contrast, according to our model, galantamine affected inhibitory control in a dose dependent fashion, reducing inhibition failures at low doses and increasing them at higher levels. These effects were sufficiently specific that the computational analysis allowed for identifying the drug administered to an individual with 70% accuracy.

**Conclusions:** Our results do not support the hypothesis that elevated tonic dopamine strongly impairs inhibitory control. Rather levodopa improved the ability to select correct actions. Instead, inhibitory control was modulated by cholinergic drugs. This approach may provide a starting point for future computational assays that differentiate neuromodulatory abnormalities in heterogeneous diseases like schizophrenia.

## Introduction

Schizophrenia is a heterogeneous clinical entity: patients with comparable symptoms show highly variable treatment responses and clinical trajectories over time (1; 2). A key challenge is to devise procedures for differential diagnostics that disambiguate potential disease mechanisms and inform individualized treatment (3). One proposal derives from the “dysconnection hypothesis” which posits that the schizophrenia spectrum consists of different abnormalities in dopaminergic and cholinergic modulation of NMDA receptor dependent plasticity (4–6). This suggests the development of assays of neuromodulation that can operate on individualized clinical data.

Eye movements are attractive targets in this regard (7). They (*i*) can be easily measured in clinical settings, (*ii*) are sensitive to changes in neuromodulation, and (*iii*) display abnormalities in schizophrenia. Saliently, it has been consistently reported that patients with schizophrenia make more errors than control participants in the antisaccade task (8–11). In this paradigm, subjects are required to saccade in the opposite direction of a visual cue. This is assumed to probe participants’ ability to inhibit a reflexive (pro)saccade towards the cue and to select and initiate the correct action, i.e., an (anti)saccade in the opposite direction (8). However, it remains unclear whether the elevated error rate (ER) in schizophrenia is caused by deficits in inhibitory control of reflexive prosaccades, in selecting correct actions (antisaccades), or by a combination of these factors.

All of these options are thought to be related to abnormal neuromodulation. Specifically, aberrant tonic dopamine (DA) levels in the basal ganglia (BG) could lead to abnormalities in the ‘NO GO’ pathway responsible for the inhibition of reflexive saccades (9–12). However, other DA-dependent mechanisms are conceivable. For example, the findings that (a) lesions in the BG do not affect antisaccade performance (13), but (b) prefrontal lesions critically impair it (14; 15), challenge the view that higher ER in schizophrenia is caused exclusively by impaired inhibitory control (16; 17). Instead, higher ER may be caused by DA-dependent processes related to selecting the correct action, e.g., aberrant prefrontal task set maintenance (17).

In contrast to the conjectured effect of elevated basal tonic DA, pro-cholinergic drugs targeting nicotinic receptors have been postulated as possible treatments for negative symptoms and cognitive impairments in schizophrenia (18–20). While results from clinical studies have been mixed (21–24), several studies have specifically investigated whether nicotine impacts antisaccade performance (25–34). These reports indicate that nicotine reduces ER (26; 29; 30; 33; 35) although see (34).

Muscarinic receptors might also be important for the antisaccade task. Indeed, the BG are rich in muscarinic receptors and receive strong cholinergic projections (36). Moreover, ACh has been suggested to play a role in the inhibition of reflexive actions towards salient stimuli (37). According to this theory, cholinergic interneurons in the striatum transiently enhance the response of the ‘NO GO’ pathway when a stimulus is suddenly presented. Thus, it is plausible that ACh regulates the inhibition of reflexive saccades during the antisaccade task.

In summary, the effects of pro-cholinergic and pro-dopaminergic drugs on the antisaccade task are not fully understood. The goal of the present study was twofold. First, we investigated the effects of pro-dopaminergic and pro-cholinergic drugs (levodopa/galantamine) on inhibitory control and action selection in the antisaccade task. Second, we asked whether these effects were specific enough to infer, based on computational modeling of antisaccade performance, which drug had been administered to a given subject. This would establish the plausibility of an assay of dopaminergic and cholinergic neuromodulation based on the antisaccade task.

To address these questions, we performed two twin experiments following a double-blind, placebo-controlled, between-group design. To uncover the effects of levodopa and galantamine on antisaccades, we used the *Stochastic Early Reaction, Inhibition and late Action* (SERIA) model (38–40), a recent computational model of antisaccade mechanisms that quantifies the contribution of inhibitory control and action selection to ER and RT. In addition, we investigated whether the parameters inferred by the model were predictive of the drug administered to individual participants. For this, we combined SERIA with a machine learning algorithm to predict the drug applied on a subject-by-subject basis. A successful prediction would speak to the translational potential of SERIA as a computational assay of dopaminergic and cholinergic neuromodulation (41).

## Methods

### Experiment and Apparatus

All procedures described here were approved by the Cantonal Ethics Committee Zurich (KEK-ZH-Nr.2014-0246). The placebo data from Exp. 1 were used in a previous study (42).

#### Participants

Participants were approached through the recruitment system of the University of Zurich. During the first visit, subjects provided written informed consent, and medical and demographic information. Only male participants were recruited due to interactions between the menstrual cycle and dopaminergic medication (43). Subjects who fulfilled all inclusion criteria (cf. Supp. 1) were invited to two experimental sessions separated by one to eight weeks.

#### Pharmacology

Two drugs were used: levodopa and galantamine. Levodopa is a precursor of DA that crosses the blood-brain barrier and increases the presynaptic availability of DA (44). Galantamine is an acetylcholinesterase inhibitor, that increases the availability of ACh at the synaptic cleft, and an allosteric potentiating ligand of the α7 (45) and αβ2 ACh nicotinic receptors (46–48).

#### Experimental procedure

At the beginning of each session, participants were orally administered color and shape matched capsules containing either Madopar® DR 250g (200mg levodopa, 50mg benserazide) or lactose (Exp. 1), or Reminyl® (8mg galantamine) or lactose (Exp. 2). Both experimenters and participants were blind to the drug administered. Subsequently, participants received written instructions regarding the experiment and participated in a training that lasted 20 to 30 minutes.

Testing started 70 minutes after drug administration. This delay was chosen to allowed both compounds to reach peak plasma levels (Madopar: 0.7h (49), Reminyl: 0.8-2h (50)). Furthermore, the half-life of levodopa is close to 1.5h (49), whereas galantamine’s half-life is 5.2h (50), and thus much longer than the mean duration of the experiment (30min).

#### Task design

Fig. 1A depicts the task procedure. A complete description can be found in (42) and in Supp. 2.

**Figure 1:**
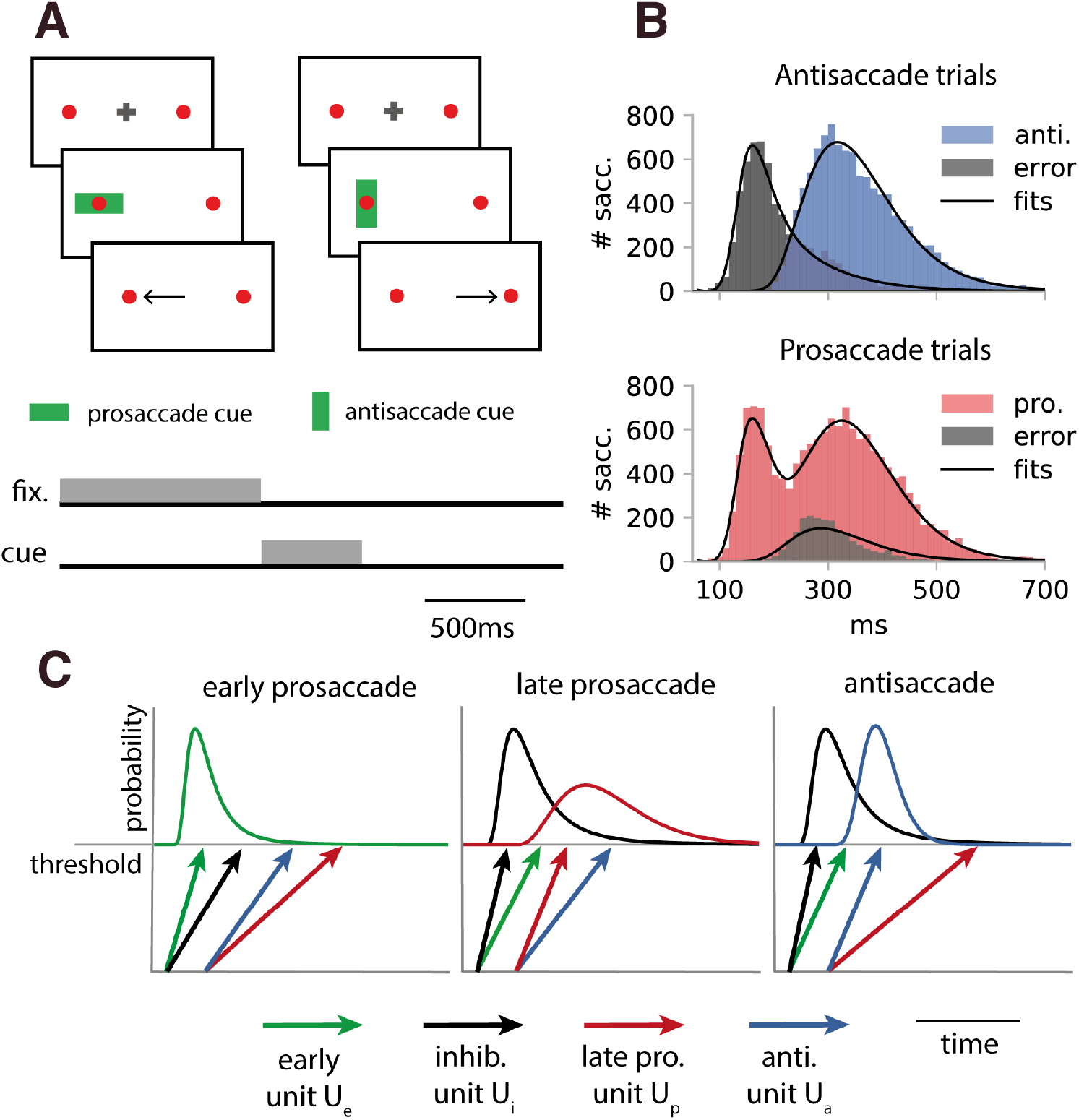
**A.** Task design: Two red circles were presented at 12° to the right and left of the center of the screen. Each trial started with a fixation cross for 500 to 1000ms. After the fixation period, a green bar was displayed for 500ms centered on one of two peripheral red circles. Participants were required to saccade to a cued red circle (prosaccade trials) or to saccade in the uncued direction (antisaccade trials) as fast as possible. **B.** RT histogram and model fits of all subjects in a subset of the data (PP50 condition). Top panel: Antisaccade trials. Correct antisaccades are displayed in blue. Errors in gray. Bottom panel: Prosaccade trials. Correct prosaccades are displayed in red. Errors in gray. Note that prosaccades were bimodally distributed. The first peak corresponds to reflexive (early) prosaccades, whereas the second peak corresponds to voluntary (late) prosaccades. The RT distributions predicted by the model are displayed in black. **C.** Schematic presentation of the model. SERIA uses four race-to-threshold units. The early unit (green) triggers fast prosaccades. If the inhibitory unit (black) hits threshold before the early unit, voluntary prosaccades (red) or antisaccades (blue) can be generated. Modified with permission from (39).

The main experiment consisted of three blocks of 192 trials. Every block contained randomly interleaved pro- and antisaccade trials, of which 20, 50 or 80% were prosaccade trials (conditions PP20, PP50, PP80 respectively). The order of the blocks was identical in both sessions, but pseudo-randomized across subjects.

### Modeling

The first main goal of this study was to quantify the effects of levodopa and galantamine on inhibitory control and action selection. The key observation here is that to complete an antisaccade, two things need to happen. First, a reflexive saccade to the peripheral cue must be stopped. Second, participants need to apply the rule associated with the cue (*vertical bar = antisaccade*) to select the corresponding action (a saccade in the direction opposite to the cue). These steps allow for different types of error: either a reflexive prosaccade is not stopped (an inhibition failure), or the wrong action is selected (a choice error).

In the case of correct prosaccades, a similar process takes place with an important twist: inhibition failures are correct responses on prosaccade trials. However, when reflexive saccades are stopped, subjects still need to select the correct action associated with the cue (*horizontal bar = prosaccade*). When the wrong action is selected, an (error) antisaccade is generated.

To quantify the effects of levodopa and galantamine on inhibitory control and action selection, it is therefore necessary to disentangle when subjects fail to inhibit reflexive prosaccades (inhibition failures), and when they fail to select the correct action (choice errors). Because none of these can be directly measured, we fitted the SERIA model to individual RT distributions (Fig. 1B-C and Supp 3).

In brief, SERIA asserts that saccades are the result of the competition between four race-to-threshold processes or units (see Fig. 1C and (39; 42)): an early response unit *U*_*e*_ associated with fast prosaccades, an inhibitory unit *U*_*i*_ whose function is to stop fast prosaccades, and two late response units that represent voluntary (late) prosaccades (*U*_*p*_) and antisaccades (*U*_*a*_). Conceptually, SERIA postulates that early or reflexive prosaccades are generated when the early unit is not stopped by the inhibitory unit. In addition, when a fast prosaccade is stopped, a voluntary eye movement is generated. The action selected (antisaccade or late prosaccade) is determined by the late unit that hits threshold first.

The model can be used to infer on several quantities that are not directly measurable. First, SERIA’s parameters capture the probability of an inhibition failure, i.e., the probability that the early unit hits threshold before all other units. Second, it is possible to quantify the mean hit time of the late units. For antisaccades, this quantity is similar to the mean RT. For prosaccades, this quantity represents the RT of voluntary (late) prosaccades. Finally, the parameters of the model determine the probability of choice errors. On an antisaccade trial, a choice error is a voluntary prosaccade. By contrast, on a prosaccade trial, an error antisaccade is generated when the antisaccade unit hits threshold before the late prosaccade unit.

Details about the modeling approach and fitting procedure can be found in Supp 3.

#### Statistical analysis

Statistical analyses were conducted using generalized linear mixed effect models (GLME) implemented in *R* (3.4.3). Subjects were entered as a random effect, whereas the factors *switch trial* (SWITCH), *prosaccade trial probability* (PP), SESSION, DRUG (drug vs. placebo), *experiment* (EXP, dopamine vs. galantamine), and DOSE defined as *drug(mg)/weight(kg)*, were treated as fixed effects. In addition, we considered the following interactions PP*SESSION, PP*SWITCH, PP*DRUG, DOSE*DRUG. When both experiments were analyzed together, we also included the interactions DRUG*EXP, DRUG*DOSE*EXP and DRUG*PP*EXP. ER were analyzed with binomial regression models, whereas probabilities were analyzed with Beta regression models. Statistical inference about RT were based on *F-*tests. For ER and probabilities, Wald tests were employed. Significance was asserted at *α* = 0.05.

#### Classification

The second main goal of this study was to test whether it is possible to determine if a given participant received levodopa or galantamine based on computational parameters derived from our model. To this end, a supervised classification algorithm was trained on individual model-based features computed from parameter estimates, with the aim to predict the drug administered on a subject-by-subject basis. More concretely, the features used to train the classifier were subject-specific differences in parameter estimates between the drug and placebo conditions. This “generative embedding” (51) strategy is a way to enhance (un)supervised learning by using posterior estimates from a generative model, instead of raw data, as a denoised and low-dimensional feature space. Classification was performed using *gradient boosting* (52) implemented in *xgboost* (53). The details of the classification strategy are explained in Supp. 4.

## Results

### Participants

In Exp. 1, 46 subjects (mean age 23.6, std. 2.9) were included in the final analysis. In Exp. 2, 44 subjects were included in the final analysis (mean age 22.4, std. 2.3). For further details see Supp. 5.

### Error rate and reaction time

Before analyzing the behavioral data with SERIA, we report the empirical ER and RT. The former is assumed to measure participants’ ability to stop reflexive prosaccades, and therefore elevated ER is thought to reflect poor inhibitory control (8). There is less consensus on what changes in RT may indicate (56; 57). For example, RT is thought to represent attentional shifting velocity (54) and saccadic processing velocity (58). An extended overview of behavioral effects is presented in Supp. 6 and 7.

#### Error rate

The mean ER on pro- and antisaccade trials is displayed in Fig. 2 top row. High congruent trial type probability was associated with fewer errors on pro- (*p* < 10^−5^) and antisaccade trials (*p* < 10^−5^). For example, participants made fewer prosaccade errors (antisaccades) when prosaccade trials were most common (PP80 block), compared to other blocks (PP20 and PP50).

**Figure 2:**
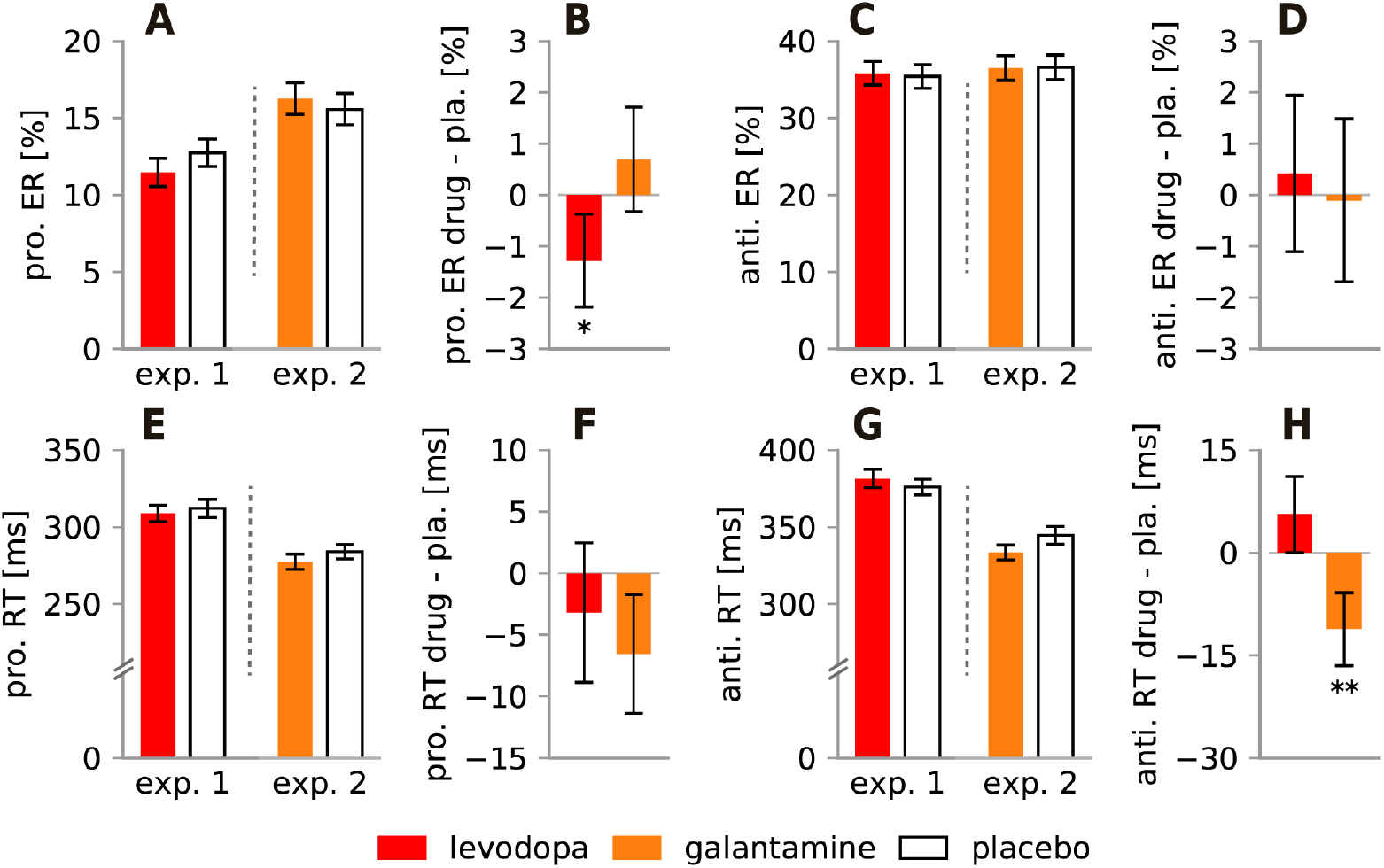
**A.** Mean prosaccade ER in Exp. 1 and 2. **B.** Difference in prosaccade ER between the drug and placebo conditions. Levodopa significantly reduced prosaccade ER (*p* = 0.010). **C.** Mean antisaccade ER. **D.** Difference in antisaccade ER between the drug and placebo conditions. **E.** Mean prosaccade RT in Exp. 1 and 2. **F.** Difference in prosaccade RT between the drug and placebo conditions. **G.** Mean antisaccade RT. **H.** Difference in antisaccade RT between the drug and placebo conditions. Galantamine decreased antisaccade RT (*p* < 10^−3^). Error bars depict the SEM. PP: prosaccade trial probability; * : *p* < 0.05; **: *p* < 0.01.

#### Error rate - drug effects

Levodopa reduced the ER on prosaccade trials (*p* = 0.010). This effect was dose dependent (*p* = 0.004). On antisaccade trials, we found no significant main effect of DRUG in Exp. 1 or 2, but there was a significant interaction between DRUG and DOSE in Exp. 2 (*p* < 10^−5^). Galantamine increased antisaccade ER at high doses, while it reduced it at more moderate levels.

#### Reaction time

RT on correct trials were analyzed similarly to ER (Fig. 2 bottom row). Higher congruent trial type probability led to lower RT in both pro- (*p* < 10^−5^) and antisaccade trials (*p* < 10^−5^).

#### Reaction times - drug effects

Levodopa increased the latency of antisaccades compared to galantamine (Fig. 2H; *p* < 10^−3^). When the two experiments were analyzed independently, we found that galantamine decreased antisaccade RT (*p* < 10^−3^). No other effect was significant.

### Modeling

The classical behavioral analysis revealed three drug related effects: (*i*) Levodopa reduced the ER on prosaccade trials, (*ii*) galantamine reduced antisaccade latency and (*iii*) increased the antisaccade ER in a dose dependent fashion. In order to relate the behavioral findings to inhibitory control or action selection, we applied computational modeling to our behavioral data. Our main goal was to determine whether levodopa and galantamine affected (*i*) the hit time of the inhibitory and late units, (*ii*) the probability of inhibition failures (inhibitory control), and (*iii*) the probability of choice errors (action selection). Drug effects on the hit times of the inhibitory or late units would demonstrate effects specific to either inhibitory control or action selection.

#### Threshold hit times

The hit times of the inhibitory, and late pro- and antisaccade units were analyzed as in the previous section. Contrary to raw RT, these can be imputed directly to the inhibition of reflexive prosaccades or to voluntary actions. For the late units, we report the expected hit times on correct trials. In the case of the inhibitory unit, pro- and antisaccade trials were analyzed together.

In agreement with (39; 42), we found that the hit time of the inhibitory unit increased with the frequency of prosaccade trials (*p* < 10^−3^), indicating reduced inhibition when prosaccade trials were more common.

#### Threshold hit times – Drug effects

Levodopa increased the latency of voluntary actions (Fig. 3A-B; late pro. *p* = 0.004; anti. *p* = 0.010). On average, voluntary saccades were 5ms slower under levodopa than under placebo, which correspond to small effect sizes (Cohen’s *f*^2^ *pro*. : 1.29, *anti*. : 1.11). However, although similar in magnitude, the effect of levodopa on the inhibitory unit failed to reach significance (*drug* − *placebo* = 5*ms*, *f*^2^ = 0.07, *p* = 0.079).

**Figure 3:**
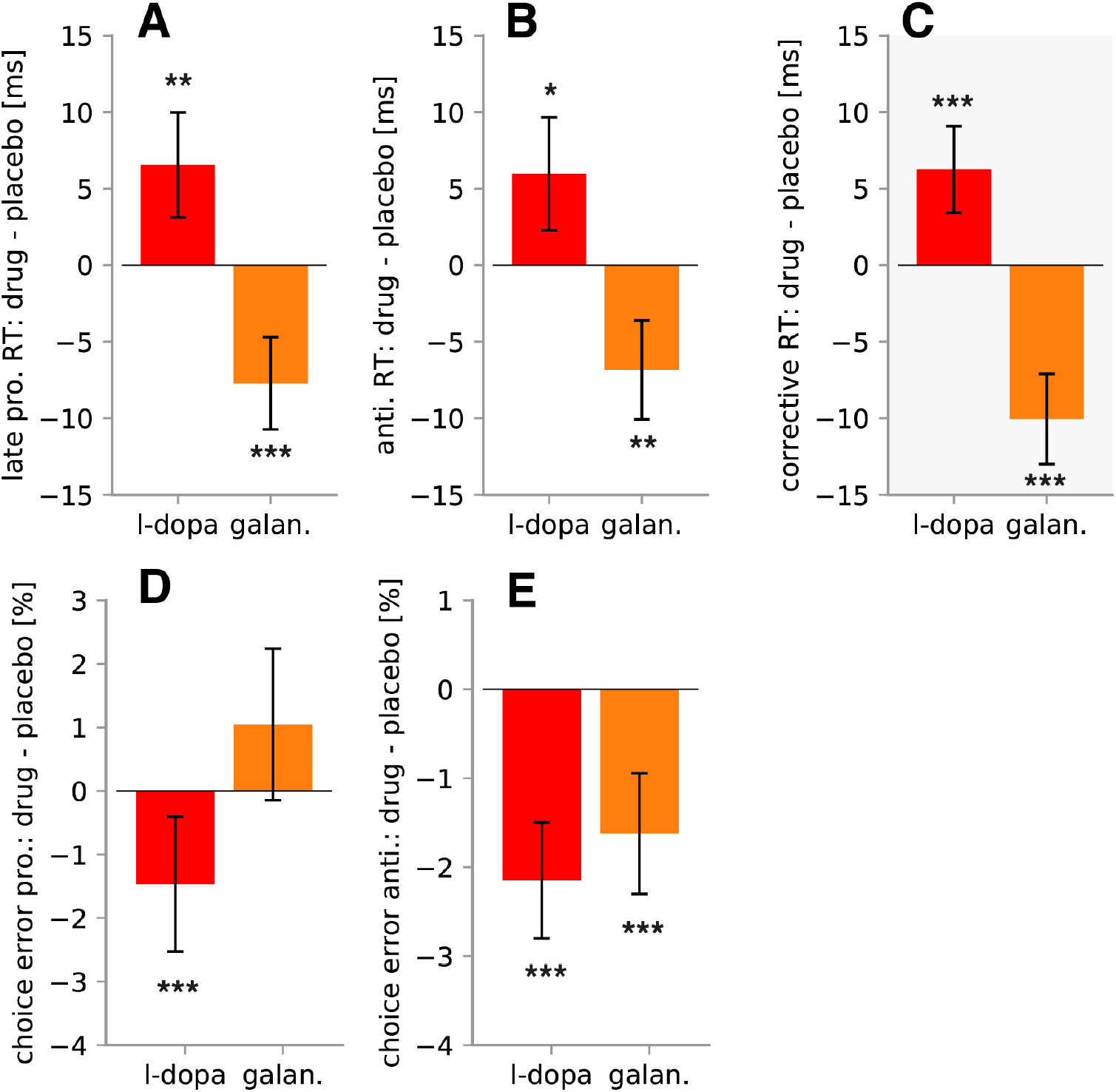
**A.** Difference in RT of late prosaccades between drug and placebo conditions (*p* < 10^−3^). **B.** Difference in RT of antisaccades (*p* < 10^−3^). **C.** Difference in RT of corrective antisaccades (*p* < 10^−3^). Note: The data in C (in gray) are completely independent of the modeling that led to the results in A and B. Corrective antisaccades display the same drug effects as voluntary saccades. **D.** Difference in ER in late prosaccades between the drug and placebo conditions (*p* < 10^−5^). **E.** Difference in late ER on antisaccade trials (*p* < 10^−3^). Error bars represent the SEM. *: *p* < 0.05; **: *p* < 0.01, ***: *p* < 0.001.

Galantamine had the opposite effect of levodopa on voluntary actions. Specifically, it reduced the hit time of late pro- (*p* < 10^−3^) and antisaccades (*p* = 0.001). On average, the hit times were 6ms lower under galantamine compared to placebo, which constitute medium effect sizes (*pro*. : *f*^2^ = 1.82; *anti*. *f*^2^ = 1.52). Again, there was no main effect of DRUG on the inhibitory unit (*p* = 0.382) but there was a dose dependent effect as explained later on.

#### Corrective antisaccades

In (39; 42), we showed that corrective antisaccades that follow errors on antisaccade trials are distributed like late responses up to a fixed delay. Consequently, SERIA predicts that corrective antisaccades should display the same drug effects as antisaccades, i.e., slower corrective antisaccades in the levodopa condition and faster antisaccades in the galantamine condition.

We analyzed 5696 corrective saccades in Exp. 1 and 4996 in Exp. 2. Because the frequency of corrective antisaccades varied widely over subjects and conditions, we accounted for the inhomogeneous number of trials by analyzing trial-by-trial RT as opposed to mean RT, using a strategy similar to (55).

Supporting our hypothesis (Fig. 3C), levodopa increased the RT of corrective antisaccades (Δ*RT* = 8*ms*; *f*^2^ = 1.11; *p* < 10^−3^), whereas galantamine had the opposite effect (Δ*R T* = −10*ms*; *f*^2^ = 1.52; *p* < 10^−3^).

#### Inhibition failures and choice errors

We proceeded to investigate the probability of choice errors and inhibition failures. Choice errors occur when the incongruent voluntary action hits threshold before the congruent action. In other words, choice errors happen when the wrong voluntary action is selected. An inhibition failure occurs when the early unit hits threshold before all other units.

Choice error rate was anticorrelated with congruent trial type probability (late pro: *p* < 10^−5^; anti: *p* < 10^−5^). Thus, the correct voluntary action was selected most often when the probability of the corresponding trial type was the highest. The probability of an inhibition failure was positively correlated with prosaccade trial probability (*p* < 10^−5^). This indicates that inhibitory control was released as prosaccade trials became more common.

#### Inhibition failures and choice errors – drug effects

Levodopa significantly reduced the probability of choice errors on pro- and antisaccade trials (Fig. 3D-E; pro.: Δ = 1.5%; *p* < 10^−3^; anti.: Δ = 2.1%; *p* < 10^−5^). By contrast, levodopa increased the probability of inhibition failures, although this effect was not significant (Δ = 1.6%, *p* = 0.082). Therefore, levodopa mainly improved the ability to select correct voluntary actions, at the cost of higher RT.

Galantamine decreased the probability of choice errors on antisaccade trials (*p* < 10^−5^). On prosaccade trials, galantamine did not have a significant effect (*p* = 0.095). There was no significant main effect of galantamine on the number of inhibition failures (*p* = 0.590).

#### Dose dependent effects

In addition to the main effects of galantamine and levodopa, we investigated any dose dependent effect. At low doses, levodopa reduced the probability of choice errors on prosaccade trials. This effect was reversed at higher doses (Fig. 4A; *p* < 10^−5^). While the main effect of DRUG was not significant in Exp. 2, galantamine had a highly significant dose dependent effect on the latency of the inhibitory unit (Fig. 4B; *p* < 10^−3^). This was reflected by a linear effect on inhibition failure probability (*p* < 10^−3^). At low doses, galantamine reduced the hit time of the inhibitory unit and the inhibition failure probability, and this effect was reversed at higher doses.

**Figure 4:**
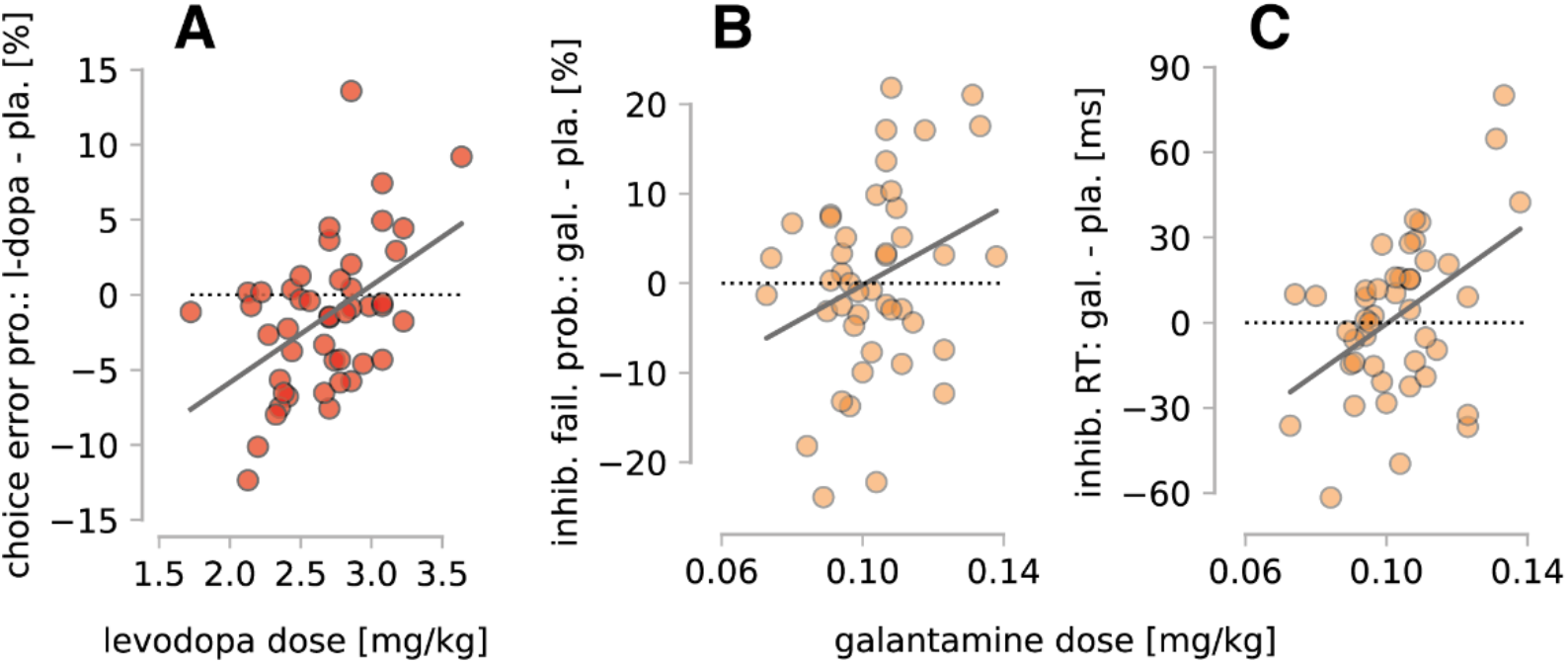
Dose dependent effects. **A.** Difference (*levodopa - placebo*) in choice ER on prosaccade trials as a function of dose in Exp. 1 (*p* < 10^−5^). At a high dose, levodopa increased the number of errors, whereas at more moderate levels, it had the opposite effect. **B.** Difference (*galantamine - placebo*) in the percentage of inhibition failures averaged across conditions. Galantamine increased the number of inhibition failures as a function of dose (*p* < 10^−3^). **C.** Difference (*galantamine - placebo*) in the RT of the inhibitory unit averaged across conditions. Galantamine increased the latency of the inhibitory unit as a function of dose (*p* < 10^−3^).

#### Classification of drug effects

Finally, we tested whether the effects of levodopa and galantamine on computational parameters can be used to predict which of the two drugs was administered (Fig. 5). Leave-one-out cross-validation resulted in 70% predictive accuracy (95% CI [61%, 79%]). A permutation test, in which the levodopa and galantamine labels were randomly swapped, showed that the predictive accuracy was highly significant (*p<0.001*). A second permutation test (in which the drug and placebo labels were randomly swapped) yielded a similar result (*p=0.001*; Fig. 5). Because drug/placebo labels (but not experiment labels) were permuted, this second test rules out that the accuracy of the classifier depended on a difference between experiments not related to the drug administered.

**Figure 5:**
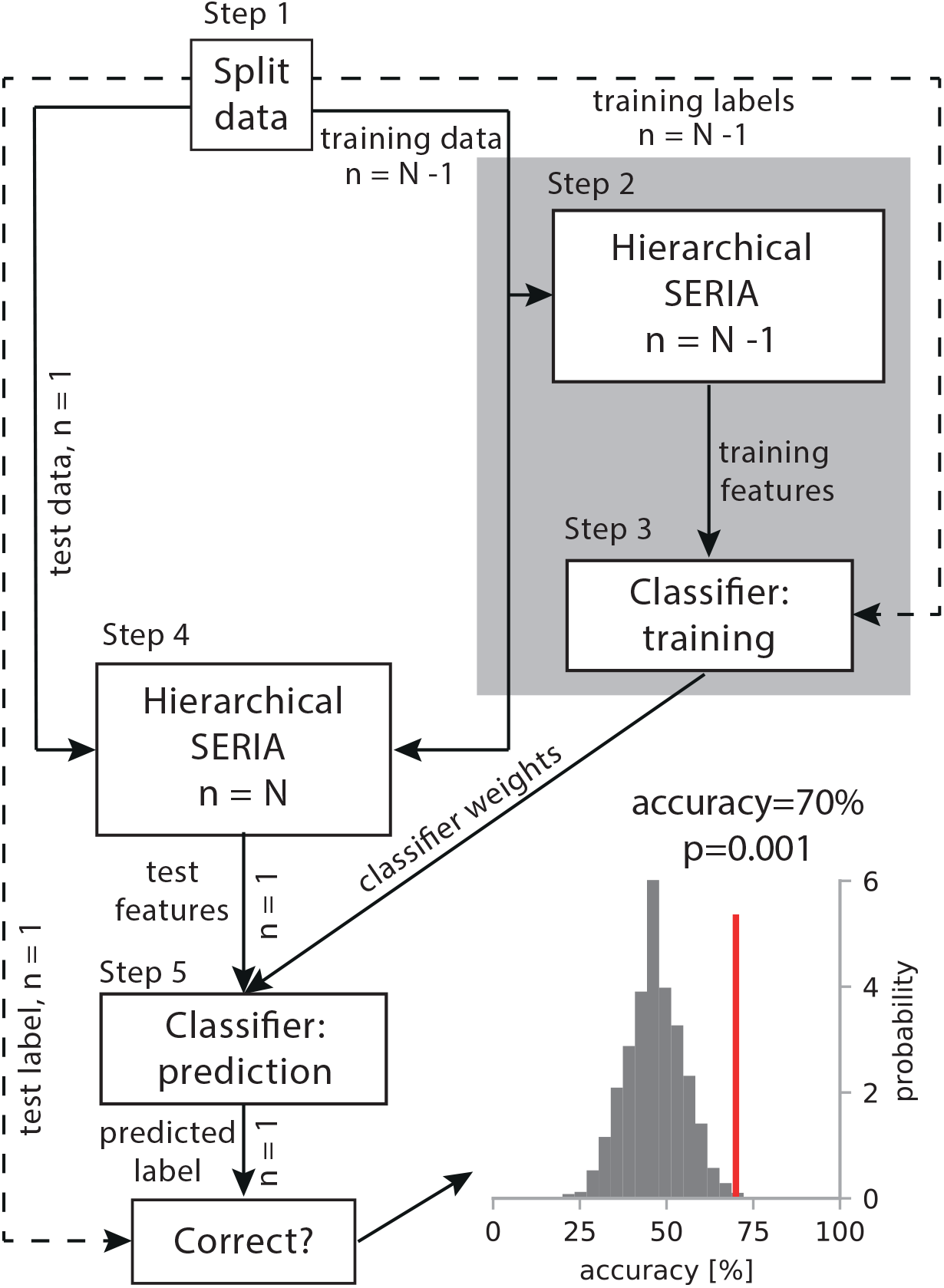
Prediction of drug labels with SERIA. This figure summarizes the classification procedure. **Step 1.** Data from N=90 subjects were split into test and training sets leaving one subject out at each iteration. **Step 2.** To generate training features, the SERIA model was fitted to data from N-1 subjects. **Step 3.** A gradient boosting classifier was trained on the SERIA parameter estimates using the drug labels from the previous step. **Step 4.** Test features were generated by fitting SERIA to data from all N subjects. **Step 5.** Weights from classifiers trained on N-1 subjects were used to predict the drug label of the left-out subjects. This resulted in a predictive accuracy of 70% (95% CI[61,79]), *p=0.001*. The histogram in the bottom right shows the accuracies using randomly permuted drug labels. The red line illustrates the true accuracy. This splitting of the data assured that the test data could not influence the training of the classifier in any way.

## Discussion

This study was motivated by the longstanding observation that aberrant neuromodulatory signaling might underlie the pathophysiology in schizophrenia (56–59). Hence, non-invasive readouts of neuromodulatory processes in patients might be of clinical relevance (4). A first test of the feasibility of such readouts can be obtained from pharmacological studies in healthy volunteers using a paradigm with consistently altered behavior in schizophrenia and with hypothesized links to potential changes in neuromodulatory transmission. The antisaccade paradigm fulfills these criteria. We have thus studied changes of its key cognitive subcomponents – in particular, inhibitory control and action selection – under pharmacological manipulations of DA and ACh.

With this goal in mind, we investigated the effect of a pro-dopaminergic (levodopa) and a pro-cholinergic (galantamine) drug on inhibition and action selection during the antisaccade task. Traditional behavioral metrics revealed several significant effects of these drugs. A more fine-grained analysis was possible through computational modeling, which indicated that levodopa altered action selection. Levodopa also increased the number of inhibition failures, although this effect was not significant. In other words, levodopa mainly enhanced the decision process between competing voluntary actions, without reliably affecting the inhibition of reflexive saccades. Higher action selection accuracy came at the cost of higher RT.

Galantamine affected both action selection and inhibitory control. Specifically, voluntary actions were facilitated by galantamine: RT were lower compared to placebo. Galantamine also improved the inhibition of reflexive actions at lower doses but had the opposite effect at higher levels. Thus, contrary to commonly held hypotheses (9; 10), dopaminergic neuromodulation affected action selection rather than inhibitory control. However, cholinergic neuromodulation strongly affected inhibitory control. Notably, these effects were specific enough to allow for identifying the administered drug on a subject-by-subject basis with reasonable accuracy. This suggests the potential for a future translation of our method into clinical applications.

In the following, we discuss our findings in relation to levodopa, galantamine, and possible clinical applications.

### Effects of levodopa

Although levodopa has been used widely in translational research (60), it has not been studied systematically in the antisaccade task (but see (61; 62)). Nevertheless, it has been hypothesized that increased tonic DA levels in the BG impair inhibitory control, which should explain the deficits observed in schizophrenia (9; 10).

According to SERIA, levodopa did not significantly alter the inhibition of reflexive saccades. However, there was a trend towards more inhibition failures in the levodopa condition. Previous studies have also failed to find changes in sto-signal RT under levodopa compared to placebo (63; 64), suggesting that increased tonic DA might have a limited effect on response inhibition.

Intriguingly, modeling demonstrated that levodopa influenced action selection in two ways: It reduced the probability of errors in selecting voluntary actions (choice errors), and it increased the latency of this type of actions. These effects were not restricted to antisaccades but extended to voluntary prosaccades and to corrective antisaccades. Thus, the effects of levodopa were most prominent in action selection and not in inhibitory control.

Prefrontal areas represent voluntary cue-action mappings in the antisaccade task (69; 70) and possibly implement the decision processes responsible for them (17; 65). In these regions, low-dose DA1 receptor mediated inhibition might induce stronger network stability (66) reducing (choice) ER and RT, while not affecting inhibitory control. This possibility is also supported by our finding that levodopa reduced choice errors on prosaccade trials at lower doses, while it increased the ER at higher doses (Fig. 4A), suggesting that excessive DA impairs voluntary action selection.

In summary, the main effect of levodopa was to slow down voluntary saccades, which led to fewer choice errors. From a modeling perspective, this suggests that levodopa promoted a speed/accuracy tradeoff, by increasing the latency of voluntary responses, and thus allowing for more evidence to accumulate. By contrast, there was no significant effect on the inhibition of reflexive saccades. Nevertheless, our analysis cannot rule out that DA affects inhibitory control in the antisaccade task.

### Effects of galantamine

While the effects of nicotine on antisaccades have been investigated previously (25–34; 67; 68), to our knowledge, this is the first antisaccade study applying a more general pro-cholinergic drug (as an AChE inhibitor, galantamine raises ACh levels in general). Our findings replicate previous studies in which nicotine was found to reduce antisaccade RT (25; 27; 28; 30; 33).

In addition to the effect on voluntary responses, galantamine also affected inhibition failure probability in a dose-dependent fashion. At a high dose, galantamine had a deleterious effect, whereas at more moderate levels, it improved performance. A comparable effect was reported previously (69), and it agrees with dose-dependent effects observed in vitro (48) and in vivo in rodents (70). In patients with schizophrenia, galantamine at high doses (32mg/day) impairs inhibitory control (71).

Although deficits on the antisaccade task have been related to DA dysregulation in the BG, the BG are also strongly modulated by cholinergic processes, due to local cholinergic interneurons and afferent projections from cholinergic nuclei (76). Our results suggest that cholinergic neuromodulation is also relevant to explain deficits in inhibitory control.

### Opposite effects of levodopa and galantamine: Predictive classification

One promising application of mathematical models in translational psychiatry concerns the development of *computational assays* that can generate single-subject predictions (41). Our results indicate that the effects of galantamine and levodopa could be discriminated based on SERIA parameter estimates obtained from eye movements during the antisaccade task. To our knowledge, this constitutes the first demonstration that antisaccade behavior can be used to make statements about neuromodulation in individual subjects. Because antisaccade performance can be easily measured in clinical settings and is robustly impaired in schizophrenia, the combination of SERIA with machine learning might find utility for translational applications in schizophrenia research. Specifically, if the accuracy of our approach were further increased, it could help identify clinically relevant subgroups with different abnormalities in neuromodulation, as postulated by the dysconnection hypothesis of schizophrenia (4–6). If successful, a computational assay of this sort might eventually contribute to procedures for differential diagnostics and aid individual treatment recommendations. The limitations and prospects of this approach need to be evaluated in future studies.

## Acknowledgements

This work was supported by the René and Susanne Braginsky Foundation (KES) and the University of Zurich (KES).

