## supplemental for "Computational dissociation of dopaminergic and cholinergic effects on action selection and inhibitory control"

**Supplementary 1 – Exclusion criteria**

Participants were excluded if any of the following criteria applied: age outside the range $18-40$, resting pulse rate outside the range $55-100bps$, abnormal electrocardiogram, regular smoking, recreational drug consumption in the past 6 months, serious mental or neurological illness, current use of psychiatric medication, use of medication that could interact with levodopa, benserazide or galantamine, and lactose intolerance.

**Supplementary 2 – Experimental procedure and data preparation**

All experiments were conducted in a dimly illuminated room. Participants sat $60cm$ in front of a computer screen. Eye movements were measured with an Eyelink 1000 (SR Research) at a sampling rate of $500Hz$. Saccades were detected with the algorithm provided by the eye-tracker manufacturer.

During the experiment, two red circles (0.25°) were displayed at a horizontal eccentricity of $\pm 12°$. A trial started when a central fixation cross was presented. Once participants started fixating the cross, it remained on the screen and had to be fixated for 500 to 1000ms, after which a green bar (3.48° × 0.8°) centered on one of the circles was displayed for $500ms$. The bar was shown in either vertical or horizontal orientation. Subjects were instructed to saccade to the circle cued by a horizontal bar (prosaccade trials) and to saccade to the non-cued circle if the bar was oriented vertically (antisaccade trials). The next trial started 1000ms after the bar was removed.

Data were preprocessed similarly as in previous studies (1; 2). Saccades were detected with the algorithm provided by the eye tracker manufacturer (SR Research – Ottawa, Canada), which implements a velocity (22deg/s) and (3600deg/s$^2$) acceleration threshold (3). Only saccades with a magnitude larger than 2 deg were analysed. Prosaccades were defined as saccades directed to the cued location. Antisaccades were defined as saccades directed opposite to the cue. As in our previous studies and similarly as in (4), prosaccades with latencies below 90ms and antisaccades with latencies below 110ms were not included in the final analysis.

**Supplementary 3 – Modeling**

This section presents the modeling in more detail. The description is based on previous publications and includes verbatim passages from the supplementary materials of (5).

To infer the model parameters of all subjects, we used the SERIA model (1). The likelihood function of SERIA is the joint probability of an action $A \in \{pro, anti\}$ and the associated RT $\in [0, \infty[$. According to this model, saccades are the product of the competition between four units that race to threshold: the early prosaccade unit $u_e$, the inhibitory unit $u_i$, the antisaccade unit $u_a$, and the late prosaccade $u_p$. Their hit times are treated as random variables ($U_e, U_i, U_a, U_p$ respectively). A prosaccade is generated when either *(i)* the early unit hits threshold before all other units, *(ii)* the late prosaccade unit hits threshold before all other units, or *(iii)* the early unit is blocked by the inhibitory unit, and the late prosaccade unit is faster than the antisaccade unit. Formally, we express these three cases with the following equation, which represents the probability $p$ of a prosaccade at time *t:*

$$\begin{gathered} p(A = pro, RT = t) = p(U_e = t)p(U_a > t)p(U_i > t)p\big(U_p > t\big) + \\ p\big(U_p = t\big)p(U_a > t)p(U_i > t)p(U_e > t) + \\ p\big(U_p = t\big)p(U_a > t)\int_0^t p(U_i = \tau)p(U_e > \tau)d\tau\,. \end{gathered} \qquad (S1)$$

An antisaccade at time $t$ occurs when the antisaccade unit hits threshold before all other units, or the inhibitory unit hits threshold before the early unit, and the antisaccade unit hits threshold before the late prosaccade unit.

$$\begin{gathered} p(A = anti, RT = t) = p(U_a = t)p\big(U_p > t\big)p(U_e > t)p(U_i > t) + \\ p(U_a = t)p\big(U_p > t\big)\int_0^t p(U_i = \tau)p(U_e > \tau)d\tau\,. \end{gathered} \qquad (S2)$$

Together, the two equations (S1 and S2) constitute the full likelihood function of the SERIA model. A more detailed treatment can be found in (1). The code of the SERIA model is openly available on https://github.com/translationalneuromodeling/tapas.

Based on previous studies (36-38), trials from each block were split according to trial type (pro- or antisaccade) and whether the preceding trial was of the same or different type (repeat and switch trials, respectively). A set of parameters describing each of the units was defined for each condition (e.g., repeat prosaccade trials).

To reduce the number of parameters of the model, we assumed that the early unit had the same parameters across all trial types, but the late units varied across all conditions. Other secondary parameters were equal across trial types but were allowed to differ between blocks and subjects. We have previously shown that this model best explains RT and ER in this version of the antisaccade task (5).

For regularization, we used a hierarchical model (6) in which the prior distribution of the parameters associated with a particular block and subject depended on the population distribution, which was simultaneously estimated from the data of all participants. In other words, we used an empirical prior that models the population distribution. Figure S1 presents the model in graphical notation following (7).

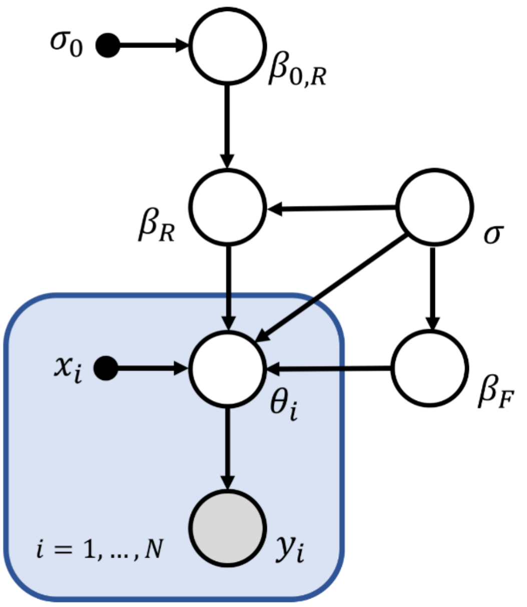


**Figure S1: Graphical representation of the hierarchical model.** Graphical summary of the statistical model using the convention in (7). Briefly, each circle represents a probability distribution and arrows indicate conditional dependence. Black dots represent fixed parameters. The equations describing the distributions of each node are given below. The most important feature of the model is that the prior distribution of each set of parameters $\boldsymbol{\theta_i}$ is parametrically defined by a set of explanatory variables $\boldsymbol{x_i}$ and coefficients $\boldsymbol{\beta}$ with variance $\boldsymbol{\sigma^2}$. These coefficients are estimated from the population distribution. We partition parameters $\boldsymbol{\beta}$ into fixed ($\boldsymbol{\beta_F}$) and random effects ($\boldsymbol{\beta_R}$), such that the latter have a prior mean estimated again from the population distribution. For the present study, random effects represent subject specific intercepts, while their mean (or global intercept) is modeled by $\boldsymbol{\beta_{0,R}}$, whose prior distribution is assumed to be centered at zero with variance $\boldsymbol{\sigma_0^2}$.

In the following, we present the mathematical description of the model depicted in Figure S1. We will write down the conditional probabilities that specify individual nodes of the graphical model, starting from the likelihood and then moving up the hierarchy. To simplify notation, we assume that the data in each run $i$, i.e., the RT and actions in a single block of each subject, are represented by a vector $y_i$ and a parameter $\theta_i$. The extension to a multivariate model is straightforward under the assumption that different parameters are conditionally independent. This assumption facilitates the construction of a hierarchical extension of SERIA.

The likelihood of the model (represented by the gray shaded circle in Figure S1) is given by the product of the likelihood of all runs, i.e. all sessions in all subjects:

$$p(y_1, \dots, y_N | \theta_1, \dots, \theta_N) = \Pi_{i=1}^{N} p(y_i | \theta_i). \qquad (S3)$$

The prior distribution of parameters $\theta_i$ is given by

$$p(\theta_i|x_i, \beta, \sigma) = \mathcal{N}(\theta_i; x_i \cdot \beta, \sigma^2). \tag{S4}$$

$X = (x_1^T, \dots, x_N^T)$ is a design matrix of size $N \times M$ that codes $M$ explanatory variables and $\beta$ is a vector of dimension $M \times 1$ that represents the effect of each explanatory variable.

In the hierarchical model, we incorporated the effect of prosaccade trial probability (levels PP20, PP50 and PP80), SESSION, and their interaction as fixed effects. The model did not include the effect of experiment or drug (verum vs. placebo), because these contain information about the labels that we eventually aimed to predict. In other words, making these factors part of the hierarchical model would have biased the results of the classifier. As a consequence, we slightly reduced the sensitivity of the hierarchical model as all parameter estimates were assumed to come from a single distribution, effectively shrinking the between-group differences. Hence, our estimate of the effect of the drugs are bound to be slightly conservative.

In addition to PP and SESSION, the factor SUBJECT was included as a random effect, that is, subject-specific intercepts were included in the design. These model the mean behavior of every participant across all conditions. The population mean was modeled as the mean of the subject-specific intercepts.

The prior distribution of $\beta$ was given by

$$p(\beta|\beta_0, \sigma) = \mathcal{N}(\beta; \beta_0, \sigma^2) \tag{S5}$$

and the prior probability of $\sigma$ was

$$p(\sigma^{-2}) = \Gamma(\sigma^{-2}; a, b). \tag{S6}$$

We distinguished between two types of $\beta$ coefficients. For fixed effects $\beta_F$, we assumed that the coefficients had a fixed prior mean $\beta_{0,F} = 0$. For random effects $\beta_R$ (in our case, the factor SUBJECT), we assumed that the prior mean $\beta_{0,R}$ is a random variable drawn from a distribution that represents the sample population, i.e. all subjects,

$$p(\beta_{0,R}|\beta_R, \sigma, \sigma_0) \propto p(\beta_R|\beta_{0,R}, \sigma)p(\beta_{0,R}|0, \sigma_0) \tag{S7}$$

$$\propto \mathcal{N}(\beta_R; \beta_{0,R}, \sigma^2)\mathcal{N}(\beta_{0,R}; 0, \sigma_0^2) \tag{S8}$$

The conditional posterior of $\beta_{0,R}$ can be computed analytically and is given by

$$p(\beta_{0,R}|\beta_R, \sigma, \sigma_0) = \mathcal{N}(\beta_{0,R}; \mu_R, \pi_R^{-1}), \tag{S9}$$

where

$$\mu_R = \frac{\sigma^{-2} \sum_{i=1}^{N} \beta_{i,R}}{N\sigma^{-2} + \sigma_0^{-2}}, \tag{S10}$$

$$\pi_R = (N\sigma^{-2} + \sigma_0^{-2}). \tag{S11}$$

The rationale for including a random effect is to account for the idiosyncrasies of each subject with the parameters $\beta_R$, while modeling a population wide intercept $\beta_{0,R}$.

All equations defined above are linear and rely on conjugate priors. Hence, it is possible to derive Gibbs steps to sample from the conditional posterior distributions of all parameters with the exception of $\theta_{1,\ldots,N}$, which are sampled from a Gaussian kernel centered at the previous sample. The model was fitted using the Metropolis-Hastings algorithm as previously described (2).

**Supplementary 4 – Classification of drug related effects**

In order to predict the drug administered to each subject, we employed gradient boosting as implemented in the package XGBoost (v0.70). Decision stumps were used as weak learners. We found that around 100 decision stumps reached the optimal performance when the learning rate was set to 0.01.

Four model-based quantities were utilized for classifying the drug labels: the mean hit time of the late pro- and antisaccade units, the mean hit time of inhibition failures and the probability of a late antisaccade. These features contain most information about inhibitory control and action selection, and we have emphasized their importance in our previous studies (2; 8).

The mean hit time of an inhibition failure is defined as

$$\frac{1}{Z}\int_0^\infty t\, p(U_e = t)p(U_i > t)p\big(U_p > t\big)p(U_a > a)dt,$$

$$Z = \int_0^\infty p(U_e = t)p(U_i > t)p\big(U_p > t\big)p(U_a > a)dt.$$

In other words, this quantifies the mean RT of early reactions that are not stopped by the inhibitory unit and reach threshold before the late units.

Only parameter estimates from the block with the highest number of antisaccade trials (80%; condition PP20) were used, as this condition was expected to be the most informative about action selection and inhibitory control. From this block, we only considered repeat antisaccades and switch prosaccades, because by design only very few trials could be switch antisaccades or repeat prosaccades. Hence, 8 features per subject entered the classifier: The four model-based quantities explained above from repeat antisaccades as well as from switch prosaccades. The classifier was trained on the *differences* between the drug and placebo conditions, effectively relying on changes between conditions to classify drugs.

Accuracy was assessed using leave-one-out cross validation. Briefly, each cross-fold consisted of the following steps (see Fig. 5). First, the data of one (test) subject was set apart. A hierarchical SERIA model was fitted to the data from the remaining N-1 subjects. Then, the classifier was trained on the difference between the parameter estimates from the drug and placebo conditions. Next, we computed the features of the test subject. To ensure similar regularization in the training and test phases, we included all N subjects in a hierarchical model, and used the parameter estimates of each subject as predictors of the corresponding drug label. This procedure was repeated once for each subject. We report the mean leave-one-out predictive accuracy across all subjects.

A possible concern about this approach is that the training data are used in the test phase to generate the features used by the classifier. Note, however, that this does not introduce any undue bias, as the training data are assumed to be known at this phase of the classification algorithm. Similarly, when, for example, the predictors used by a classifier are mean-centered (i.e. the mean is subtracted from the training set), this transformation should also be applied to the test data (i.e. subtracting the mean of the training data from the test set). Crucially, this does not introduce any bias as long as the test data is not used to compute the mean of the training data. In summary, the training data may well be used to inform the pre-processing of the test data, whereas the test data are not allowed to have an impact on the training phase. Thus, our algorithm appropriately leverages the regularization provided by the hierarchical SERIA model, without using the test data during training.

To test statistical significance, two permutation tests were conducted. First, we randomly shuffled the drug labels (levodopa/galantamine) 5000 times and reevaluated the analysis. Second, to rule out that the results were driven by any difference between Exp. 1 and 2 not related to the drug treatment, we implemented a test in which the drug and placebo labels were randomly swapped. This represents the distribution of classification accuracies when experiment but not drug labels are used to predict the substance that was administered.

### Supplementary 5 – Descriptive statistics – Participants and valid trials

In Exp 1., four participants were excluded because their data were incomplete. No subject experienced side effects. Participants were not significantly better than chance in guessing the substance administered after the first ($p$ = 0.07) and second session ($p$ = 0.75). In total, 52993 trials were analyzed, from which 1303 trials were excluded (Supp. Table. S1) using criteria specified previously (1).

In Exp. 2, four subjects reported side effects and were excluded from the study. Two subjects were excluded because of incomplete data. Again, participants could not discriminate galantamine from placebo (first session $p$ = 0.54; second session $p$ = 0.36). 50688 trials were analyzed, from which 1416 were excluded (Supp. Table. S1).

**Table S1**: Mean number of invalid trials per subject

| | **Mean # invalid trials** | | | | | | | |
|---|---|---|---|---|---|---|---|---|
| | **Experiment 1** | | | | | | | |
| | Valid | Blink | Missing | Aborted | FE | Late | Early | NS |
| Placebo | 563(10) | 4(5) | 1(3) | 1(2) | 5(5) | 0(0) | 1(1) | 0(0) |
| L-dopa | 562(14) | 5(7) | 1(3) | 1(2) | 5(6) | 0(0) | 1(2) | 0(0) |
| | **Experiment 2** | | | | | | | |
| | Valid | Blink | Missing | Aborted | FE | Late | Early | NS |
| Placebo | 558(24) | 10(17) | 3(6) | 1(1) | 3(7) | 0(0) | 1(1) | 0(0) |
| Galan. | 562(18) | 9(14) | 1(3) | 1(2) | 3(5) | 0(0) | 0(1) | 0(0) |

std. is displayed in brackets. Late = saccades with latency above 800ms. Early = saccades with latency below 90ms. FE = fixation errors, NS = No saccade.

### Supplementary 6 – Error rate

Mean ER across experiments, conditions and trial type probabilities are presented in Table **S2**.

**Table S2:** Mean error rate

| | Error rate (%) | | | | | |
|---|---|---|---|---|---|---|
| | **Experiment 1** | | | | | |
| | Placebo | | | Levodopa | | |
| | PP20 | PP50 | PP80 | PP20 | PP50 | PP80 |
| Anti. | 23(17) | 36(22) | 52(20) | 23(16) | 36(21) | 52(24) |
| Pro. | 26(15) | 11(8) | 4(3) | 25(18) | 10(8) | 3(4) |
| | **Experiment 2** | | | | | |
| | Placebo | | | Galantamine | | |
| | PP20 | PP50 | PP80 | PP20 | PP50 | PP80 |
| Anti. | 23(17) | 36(22) | 55(19) | 22(17) | 36(21) | 53(22) |
| Pro. | 31(17) | 14(10) | 5(3) | 32(16) | 15(9) | 5(4) |

std. is displayed in brackets.

Table **S3** and **S4** display the complete statistical analysis of ER in pro- and antisaccade trials (see Fig. **S2** and **S3**). In general, subjects made fewer errors in repeat trials compared to switch trials. Similarly, the mean ER was lower in the second session compared to the first session. Although the trial type probability manipulation (PP) was significant in pro- and antisaccade trials, there was no significant interaction between it and any of the drugs administered. Moreover, there was no significant interaction between the factor SWITCH and DRUG when we investigated each experiment separately (not displayed).

**Table S3:** Statistical analysis: error rate prosaccade trials

| | Chisq | Df | Pr(>Chisq) | |
|---|---|---|---|---|
| PP | 871.7 | 2 | 0.000 | *** |
| SESSION | 13.9 | 1 | 0.000 | *** |
| SWITCH | 2331.2 | 1 | 0.000 | *** |
| DRUG | 1.1 | 1 | 0.296 | |
| DOSE | 0.1 | 1 | 0.776 | |
| EXPERIMENT | 6.2 | 1 | 0.013 | * |
| PP * SESSION | 3.4 | 2 | 0.183 | |
| PP *SWITCH | 27.0 | 2 | 0.000 | *** |
| DRUG * DOSE | 1.4 | 1 | 0.235 | |
| PP * DRUG | 0.4 | 2 | 0.827 | |
| DRUG * EXPERIMENT | 6.9 | 1 | 0.009 | ** |
| PP * EXPERIMENT | 0.5 | 2 | 0.763 | |
| DOSE * EXPERIMENT | 0.5 | 1 | 0.462 | |
| PP * DRUG * EXPERIMENT | 1.6 | 2 | 0.456 | |
| DRUG * DOSE * EXPERIMENT | 7.5 | 1 | 0.006 | ** |

*:p<0.05, **:p<0.01, ***:p<0.001

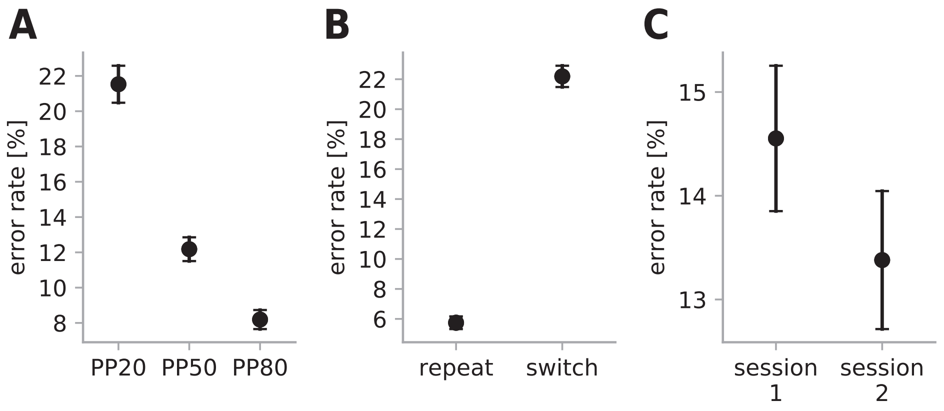


**Figure S2:** A. Mean error rate on prosaccade trials across blocks with different prosaccade trial type probability (PP). B. Mean error rate on prosaccade trials on repeat and switch trials. C. Mean error rate on prosaccade trials in session 1 and 2. Error bars display the sem..

**Table S4:** Statistical analysis: Error rate antisaccade trials

| | Chisq | Df | Pr(>Chisq) | |
|---|---|---|---|---|
| PP | 901.2 | 2 | 0.000 | *** |
| SESSION | 7.6 | 1 | 0.006 | ** |
| SWITCH | 2061.6 | 1 | 0.000 | *** |
| DRUG | 0.5 | 1 | 0.482 | |
| DOSE | 0.0 | 1 | 0.824 | |
| EXPERIMENT | 0.2 | 1 | 0.663 | |
| PP:SESSION | 8.3 | 2 | 0.016 | * |
| PP:SWITCH | 73.1 | 2 | 0.000 | *** |
| DRUG:DOSE | 13.8 | 1 | 0.000 | *** |
| DRUG:EXPERIMENT | 0.0 | 1 | 0.850 | |
| DOSE:EXPERIMENT | 0.3 | 1 | 0.564 | |
| PP:DRUG | 0.8 | 2 | 0.675 | |
| PP:EXPERIMENT | 0.2 | 2 | 0.886 | |
| DRUG:DOSE:EXPERIMENT | 11.8 | 1 | 0.001 | *** |
| PP:DRUG:EXPERIMENT | 0.5 | 2 | 0.789 | |

*:p<0.05, **:p<0.01, ***:p<0.001

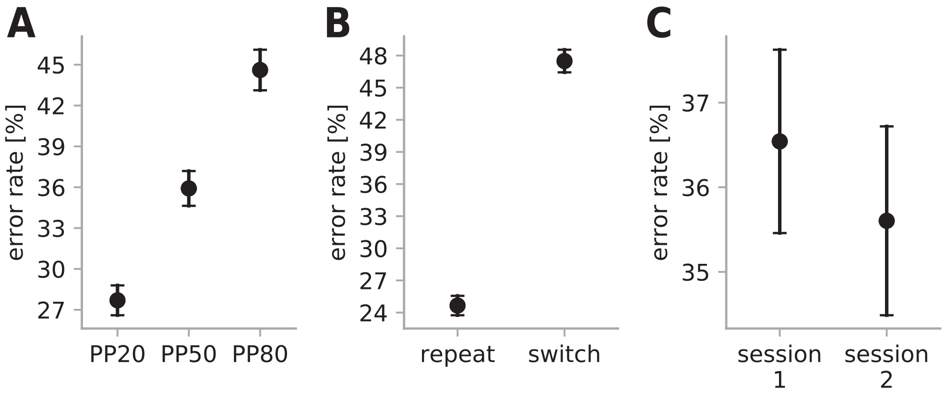


**Figure S3**: A. Mean error rate on antisaccade trials across blocks with different prosaccade trial type probability (PP). B. Mean error rate on antisaccade trials on repeat and switch trials. C. Mean error rate on antisaccade trials in session 1 and 2. Error bars display the sem..

### Supplementary 7 – Reaction times

Table **S5** presents the mean RT across trial types, trial type probabilities, experiments, and drug conditions.

**Table S5:** Mean reaction time.

| Mean reaction times (ms) | | | | | | | |
|---|---|---|---|---|---|---|---|
| | | Experiment 1 | | | | | |
| | | Placebo | | | Levodopa | | |
| Trial | Action | PP20 | PP50 | PP80 | PP20 | PP50 | PP80 |
| Anti. | Anti. | 355 | 381 | 391 | 367 | 380 | 396 |
| | | (59) | (55) | (60) | (73) | (65) | (70) |
| Anti. | Pro. | 234 | 231 | 226 | 231 | 228 | 222 |
| | | (51) | (48) | (31) | (56) | (43) | (39) |
| Pro. | Pro. | 332 | 320 | 285 | 329 | 319 | 279 |
| | | (72) | (68) | (59) | (65) | (60) | (51) |
| Pro. | Anti. | 327 | 330 | 338 | 338 | 351 | 342 |
| | | (68) | (45) | (56) | (70) | (82) | (65) |
| | | Experiment 2 | | | | | |
| | | Placebo | | | Galantamine | | |
| Trial | Action | PP20 | PP50 | PP80 | PP20 | PP50 | PP80 |
| Anti. | Anti. | 323 | 348 | 363 | 313 | 335 | 353 |
| | | (50) | (66) | (72) | (49) | (52) | (60) |
| Anti. | Pro. | 218 | 225 | 215 | 220 | 220 | 211 |
| | | (35) | (41) | (30) | (44) | (39) | (34) |
| Pro. | Pro. | 299 | 293 | 261 | 285 | 286 | 262 |
| | | (55) | (58) | (41) | (59) | (52) | (52) |
| Pro. | Anti. | 300 | 307 | 305 | 293 | 301 | 314 |
| | | (42) | (52) | (62) | (50) | (42) | (59) |

std. is displayed in brackets.

Tables **S6** and **S7** display the statistical analysis of the mean RT on pro- and antisaccade trials (see Fig. **S4** and **S5**). Similarly as with ER, saccades were faster in the second session. Regardless of trial type, correct switch trials displayed higher latencies than repeat trials. Trial type probability (PP) had a significant effect on response latency but did not interact with the drug administered.

**Table S6:** Statistical analysis of mean prosaccade reaction time.

| | **Num DF** | **Den DF** | **F value** | **Pr(>F)** | |
|---|---|---|---|---|---|
| PP | 2 | 972 | 30.4 | 0.000 | *** |
| SESSION | 1 | 972 | 31.6 | 0.000 | *** |
| SWITCH | 1 | 972 | 219.6 | 0.000 | *** |
| DRUG | 1 | 972 | 5.2 | 0.022 | * |
| DOSE | 1 | 86 | 0.8 | 0.376 | |
| EXPERIMENT | 1 | 86 | 8.4 | 0.005 | ** |
| PP:SESSION | 2 | 972 | 1.0 | 0.385 | |
| PP:SWITCH | 2 | 972 | 8.4 | 0.000 | *** |
| DRUG:DOSE | 1 | 972 | 0.4 | 0.530 | |
| DRUG:EXPERIMENT | 1 | 972 | 0.7 | 0.394 | |
| DOSE:EXPERIMENT | 1 | 86 | 0.0 | 0.961 | |
| PP:DRUG | 2 | 972 | 0.9 | 0.421 | |
| PP:EXPERIMENT | 2 | 972 | 2.0 | 0.133 | |
| DRUG:DOSE:EXPERIMENT | 1 | 972 | 3.7 | 0.054 | . |
| PP:DRUG:EXPERIMENT | 2 | 972 | 0.5 | 0.613 | |

*:p<0.05, **:p<0.01, ***:p<0.001

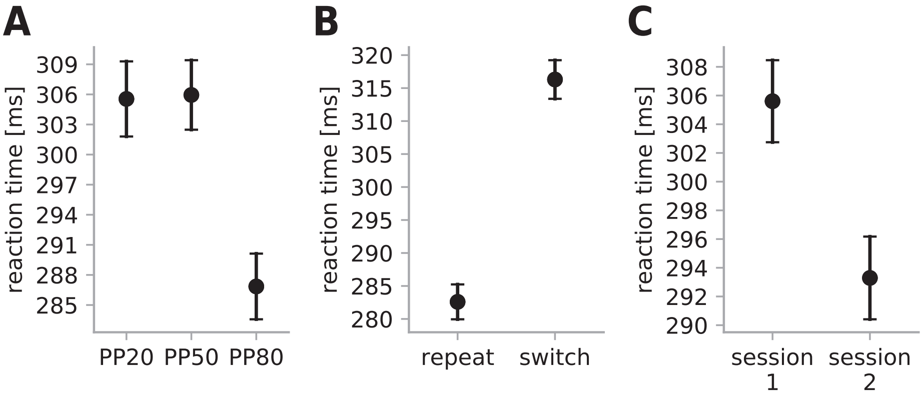


**Figure S4**: A. Mean reaction time on prosaccade trials across blocks with different prosaccade trial type probabilities (PP). B. Mean reaction time on prosaccade trials on repeat and switch trials. C. Mean reaction time on prosaccade trials in session 1 and 2. Error bars display the sem..

**Table S7:** Statistical analysis of mean antisaccade reaction time.

| | Num DF | Den DF | F value | Pr(>F) | |
|---|---|---|---|---|---|
| PP | 2 | 968.0 | 35.2 | 0.000 | *** |
| SESSION | 1 | 968.0 | 75.1 | 0.000 | *** |
| SWITCH | 1 | 968.0 | 103.3 | 0.000 | *** |
| DRUG | 1 | 968.0 | 1.5 | 0.216 | |
| DOSE | 1 | 86.0 | 0.5 | 0.504 | |
| EXPERIMENT | 1 | 86.0 | 12.6 | 0.001 | *** |
| PP:SESSION | 2 | 968.0 | 1.6 | 0.198 | |
| PP:SWITCH | 2 | 968.0 | 4.9 | 0.007 | ** |
| DRUG:DOSE | 1 | 968.0 | 2.3 | 0.128 | |
| DRUG:EXPERIMENT | 1 | 968.0 | 12.3 | 0.000 | *** |
| DOSE:EXPERIMENT | 1 | 86.0 | 0.3 | 0.602 | |
| PP:DRUG | 2 | 968.0 | 1.0 | 0.356 | |
| PP:EXPERIMENT | 2 | 968.0 | 2.4 | 0.093 | . |
| DRUG:DOSE:EXPERIMENT | 1 | 968.0 | 0.1 | 0.702 | |
| PP:DRUG:EXPERIMENT | 2 | 968.0 | 1.0 | 0.383 | |

*:p<0.05, **:p<0.01, ***:p<0.001

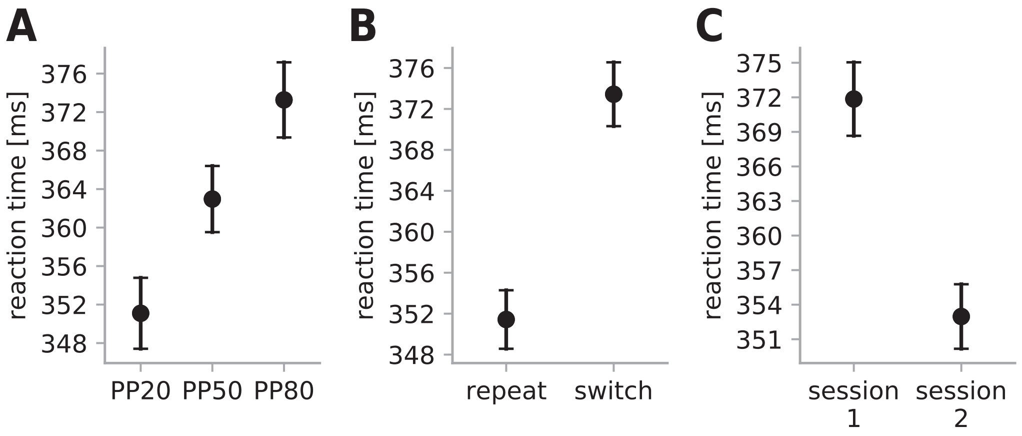


**Figure S5:** A. Mean reaction times on antisaccade trials across blocks with different prosaccade trial type probabilities (PP). B. Mean reaction times on antisaccade trials on repeat and switch trials. C. Mean reaction times on antisaccade trials in session 1 and 2. Error bars display the sem..
